# Dynamics and length distributions of microtubules with a multistep catastrophe mechanism

**DOI:** 10.1101/2022.08.19.504495

**Authors:** Felix Schwietert, Lina Heydenreich, Jan Kierfeld

## Abstract

Regarding the experimental observation that microtubule catastrophe can be described as a multistep process, we extend the Dogterom–Leibler model for dynamic instability in order to discuss the effect that such a multistep catastrophe mechanism has on the distribution of microtubule lengths in the two regimes of bounded and unbounded growth. We show that in the former case, the steady state length distribution is non-exponential and has a lighter tail if multiple steps are required to undergo a catastrophe. If rescue events are possible, we detect a maximum in the distribution, i.e., the microtubule has a most probable length greater than zero. In the regime of unbounded growth, the length distribution converges to a Gaussian distribution whose variance decreases with the number of catastrophe steps. We extend our work by applying the multistep catastrophe model to microtubules that grow against an opposing force and to microtubules that are confined between two rigid walls. We determine critical forces below which the microtubule is in the bounded regime, and show that the multistep characteristics of the length distribution are largely lost if the growth of a microtubule in the unbounded regime is restricted by a rigid wall. All results are verified by stochastic simulations.

## 1. Introduction

Assembly and disassembly of microtubules (MTs) is fundamental for many processes in the cell. The dynamics of MT polymerization is commonly described by the term *dynamic instability* [1]: a MT stochastically switches between a growing and a shrinking state in which it polymerizes or depolymerizes, respectively. The transition from growth to shrinkage is called *catastrophe*, the reverse process *rescue*. A first mathematical description of dynamic instability was provided by the Dogterom–Leibler model [2, 3], which is based on four constant parameters: a (de)polymerization velocity for the growing and the shrinking state, and two transition rates for the occurrence of catastrophes or rescues. Depending on these parameters, the MT exhibits two dynamical regimes: a regime of bounded growth with zero mean growth velocity and a stationary exponentially decaying length distribution, and a regime of unbounded growth with a non-zero mean growth velocity.

Later on, Odde *et al*. [4] and, more recently, Stepanova *et al*. [5] and Gardner *et al*. [6] found that the durations of the growth intervals are not distributed exponentially as one would expect for a constant catastrophe rate. Instead, the measured distributions could be well described by a multistep process, which means that the MT ages and the catastrophe rate increases during growth. While it was concordantly reported from control groups of in vitro experiments that a MT has to pass approximately three steps to undergo a catastrophe [4, 6, 7], it was also shown in the same experiments that the number of steps depends on concentrations of kinesins [6] or MT-targeting agents [7].

The underlying mechanism of MT aging is still under debate and several microscopic models have been proposed. For instance, it was suggested that a catastrophe is triggered by a certain number of “sub-catastrophes” of single protofilaments [8, 9, 10]. A chemomechanical approach led to the conclusion that the MT tip becomes more tapered during growth, which promotes catastrophe [11]. Another model, which included Brownian dynamics of single tubulin molecules, revealed that MT aging might be a much more complex stochastic process relying on a fluctuating MT tip and an increasing number of curled protofilaments [12]. Chemomechanical stochastic growth models on the dimer level revealed that a catastrophe could be triggered by a “nucleus” of three neighboring protofilaments shrinking by more than 6 dimers, such that its GTP-cap is removed and its ends reach into the GDP-body of the MT [13, 14].

In this paper, we do not concentrate on the microscopic details of MT aging but on the consequences that a multistep catastrophe mechanism has for the distribution of MT lengths. For that purpose, we extend the empirical Dogterom–Leibler model by subdividing the growing state into an arbitrary number *n* of sub-states a MT has to pass to undergo a catastrophe. In the bounded growth regime, the stationary form of the resulting master equations has to be solved numerically, except for the case that MTs can not be rescued. However, taking advantage of the results of Jemseena and Gopalakrishnan [15], who set up and analyzed master equations for dynamic instability with an age-dependent catastrophe rate, we are able to compute exact values for the mean MT length and higher moments and to provide an approximation that comprises the key characteristics of the length distribution for arbitrary numbers *n* of sub-states. While Jemseena and Gopalakrishnan made up heuristic functions to directly fit the age-dependency of catastrophe, our work is based on the model that catastrophe is a multistep process with equal transition rates for each step. Our main results are similar to those obtained by Jemseena and Gopalakrishnan: the duration of MT growth becomes less stochastic if more sub-steps are necessary to induce a catastrophe, which results in a more narrow length distribution with a lighter tail. In particular, the stationary distribution has a maximum if rescues are possible, i.e., the MT has a most probable length greater than zero, in contrast to the monotonically decreasing exponential distribution that follows from a single-step catastrophe. Going beyond the work of [15], we also examine the regime of unbounded growth, where the MT lengths approach a Gaussian distribution as in the case of a single-step catastrophe [2, 3] but with a variance that decreases with the number of sub-steps that are necessary to trigger a catastrophe.

Furthermore, we apply our results for multistep MT dynamics to models of MTs under an opposing force and in a rigid confinement [16, 17]. An opposing force suppresses MT growth and can counteract the effects of an increased number of required catastrophe steps *n* such that the critical force, below which the MT is in the bounded regime, becomes a monotonic function of *n*. A rigid confinement to maximum MT lengths *L* truncates the determined probability distributions to the accessible interval [0, *L*] accompanied by a finite probability to find the MT stalled at the confining boundary. Moreover, the confinement forces MTs from the unbounded regime to a stationary length distribution, which, however, does not exhibit multistep characteristics but is exponentially increasing.

## 2. Model

### 2.1. Classical single-step model

The Dogterom–Leibler model [3] is an empirical coarse-grained MT model that describes dynamic instability as stochastic switches between phases of growth and shrinkage with constant velocities. Both catastrophe and rescue are described as single-step or Poisson process as sketched in figure 1(a). The model requires four parameters: growth and shrinkage velocities *v*_*±*_ as well as catastrophe and rescue rates *ω*_c_ and *ω*_r_. The probability densities *p*_*±*_(*x, t*) of the length *x* of a growing (+) or a shrinking (−) MT obey the following Fokker–Planck equations (FPEs):

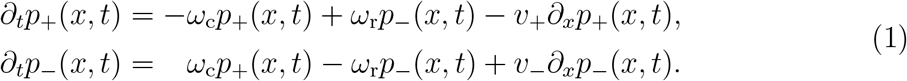

**Figure 1.**
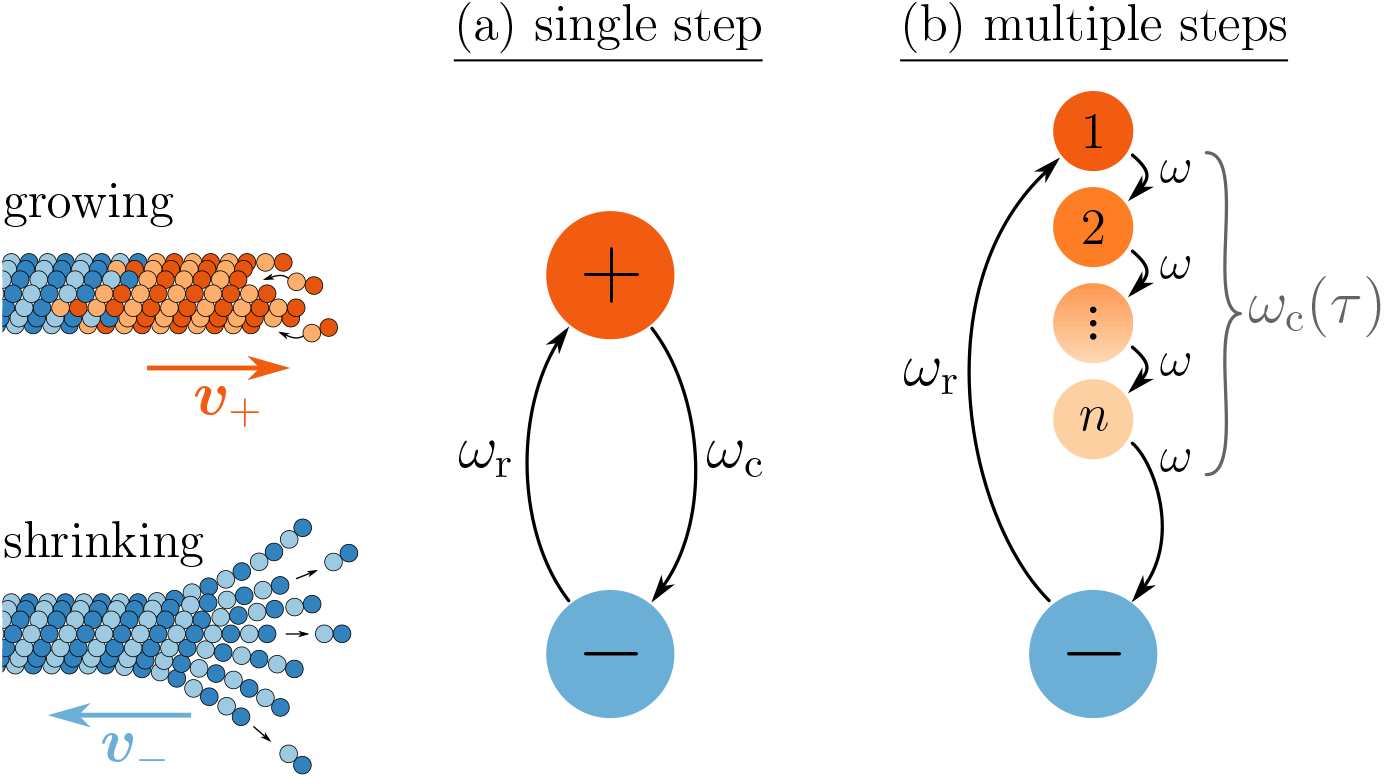
Models of dynamic instability. The MT is either in a growing or in a shrinking state, in which its tip moves with constant velocities *v*_+_ or *v*_−_. (a) In the classical Dogterom–Leibler model, catastrophe is a single step process with a constant rate *ω*_c_. (b) When catastrophe is modeled as a multistep process, the growing state is divided into *n* sub-states, and a growing MT has to pass *n* sub-steps, each with rate *ω*, before it undergoes a catastrophe. This *n*-step process can be summarized by means of an age-dependent catastrophe rate *ω*_c_(*τ*).

The mean velocity of the MT tip (time-averaged over many catastrophe and rescue cycles) is given by

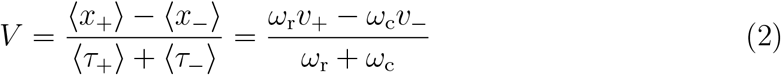

and can serve as an order parameter for the transition between bounded and unbounded regimes of MT growth. With a reflecting boundary at *x* = 0, i.e., with the MT undergoing a forced rescue as soon as it shrinks back to zero length, a negative parameter *V* < 0 leads to a zero mean velocity and bounded MT growth. Then, the overall probability density *p*(*x, t*) = *p*_+_(*x, t*) + *p*_−_(*x, t*) converges to a stationary exponential distribution with a finite mean length

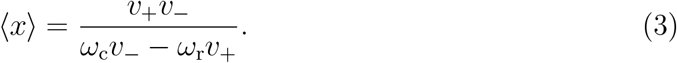

For *V* > 0, the parameter *V* is always identical to the mean growth velocity and MT growth is unbounded without a stationary length distribution. The probability density approaches a time-dependent Gaussian distribution

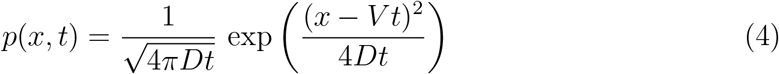

with a diffusion constant

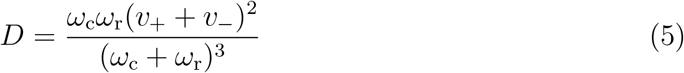

for the length diffusivity.

### 2.2. Multistep model

In the following, we extend the Dogterom–Leibler model in a way that takes account of the experimental observation that MT catastrophe is a multistep process [4, 5, 6], see figure 1(b). For this purpose, we introduce the number of steps *n* a MT has to pass to undergo a catastrophe. As long as a MT has not passed all *n* steps, it continues growing with *v*_+_. Therefore, the growing state can be divided into *n* sub-states *i* = 1…*n*. In the experiments in [4, 5, 6], the conclusion that MT catastrophe is a multistep process was derived from the observation that growth durations *τ*_+_ are gamma-distributed,

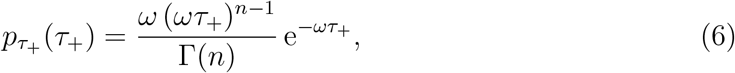

with the gamma function 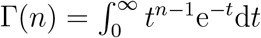 [18]. A gamma distribution implies that each catastrophe step occurs with the *same* step rate *ω* and that backward steps are not allowed, i.e., the states *i* = 1…*n* are passed in a prescribed order as sketched in figure 1(b). Fits to the experimental data gave *n* ∼ 3 [4, 6, 7] but values of *n* depend on concentrations of kinesins [6] or MT-targeting agents [7]. Since rescue is still described as a single-step process, MT dynamics is now characterized by a set of five parameters *n, v*_−_, *v*_+_, *ω*_r_ and *ω*.

Introducing *n* sub-states gives rise to a time- or age-dependent catastrophe rate via the general relation

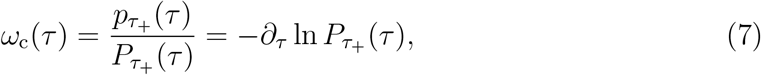

which holds for an arbitrary probability density 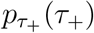 of growth durations *τ*_+_, and where 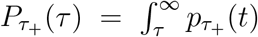 is the survival probability of a growing MT: the catastrophe rate is only observed within the ensemble of surviving MTs. For a gamma distribution (6), we have

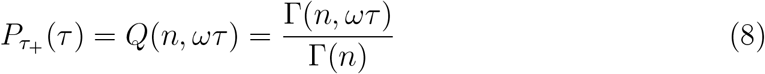

with the upper incomplete gamma function 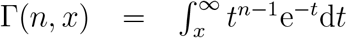 and its corresponding regularized form *Q*(*n, x*) [18]. This leads to a catastrophe rate *ω*_*c*_(*τ*), which is monotonically increasing with time *τ*. For small times *τ* ≪ *ω*^−1^, the final catastrophe is unlikely, because *n* − 1 prior sub-steps in a prescribed order are necessary that occur each with probability *ωτ* resulting in *ω*_*c*_(*τ*)*/ω* ≈ (*ωτ*)^*n*−1^*/*(*n* − 1)!. For large times *τ* ≫ *ω*^−1^, *n* − 1 prior sub-steps have passed almost certainly with probability 1 − (*n* − 1)*/ωτ*, and the rate for the final catastrophe approaches *ω*, i.e., *ω*_*c*_(*τ*)*/ω* ≈ 1 − (*n* − 1)*/ωτ*. We note that this asymptotic form of the catastrophe rate with an algebraic approach to unity is different from the exponential approach assumed by Jemseena and Gopalakrishnan [15].

With the growth duration *τ*_+_, also the length gain *x*_+_ = *v*_+_*τ*_+_ during one growth interval is gamma distributed:

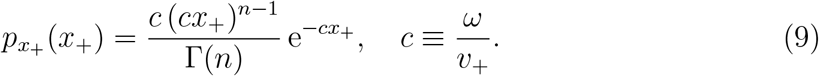

On average, a MT grows for a duration of ⟨*τ*_+_⟩ = *nω*^−1^, and its tip covers a distance of ⟨*x*_+_⟩ = *v*_+_*nω* during that interval. Two generic situations will be of interest:

i. A comparison of models that yield the same (experimentally observed) average growth duration ⟨*τ*_+_⟩ but feature a different number of sub-states *n*. Then, we have to simultaneously use a *n*-dependent catastrophe step rate *ω* ∝ *n*, i.e., if more sub-states have to be passed to catastrophe we have to increase the step rate accordingly to get the same overall average growth duration or distance.
ii. Models for experiments with microtubule associated regulating proteins, such as the catastrophe-promoting MCAK [6], where the catastrophe step rate *ω* remains fixed while the number of sub-steps *n* can be manipulated.

Together with the mean shrinking duration 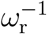 and distance 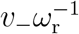, we deduce the mean tip velocity analogously to (2):

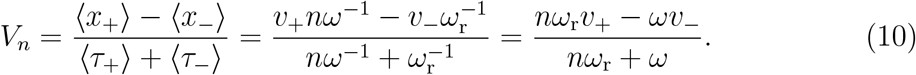

Again, the sign of *V*_*n*_ determines whether MT growth is bounded and a stationary state exists. MT growth is stabilized and may leave the bounded regime in scenario (ii) if only the number of sub-steps *n* to trigger a catastrophe is increased (*n* > *n*_c_ = *ωv*_−_*/ω*_r_*v*_+_) while the other parameters remain constant. If we compare models with the same average growth duration ⟨*τ*_+_⟩ but different number of sub-states *n* in scenario (i), on the other hand, we also have to use a catastrophe rate *ω* ∝ *n*, and *V*_*n*_ becomes independent of *n* in (10). Then, also the sign of *V*_*n*_ and, thus, the MT growth regime (bounded or unbounded) is unchanged by *n*.

For a general mathematical description, we assign a probability density *p*_*i*_(*x, t*) to each sub-state *i* = 1…*n* of a growing MT. The total growing state density is given by *p*_+_(*x, t*) = Σ_*i*_ *p*_*i*_(*x, t*). The stochastic time evolution of the probability densities is described by a system of *n* + 1 FPEs:

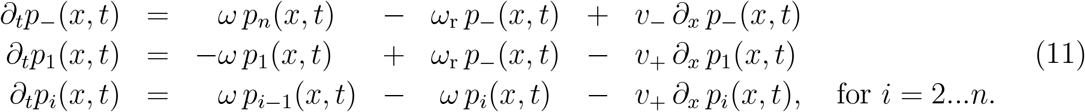

Due to the reflecting boundary, the probability current density

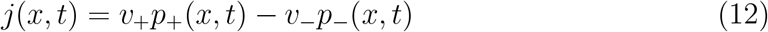

has to vanish at *x* = 0. Furthermore, in any stationary state (∂_*t*_*p*_*i*_(*x, t*) = 0), the current density is constant in space, as can be seen by summing up the FPEs (11):

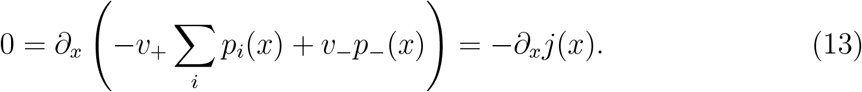

Together with *j*(*x* = 0) = 0, this implies that, in a steady state, the probability current density has to vanish everywhere. With the resulting relation

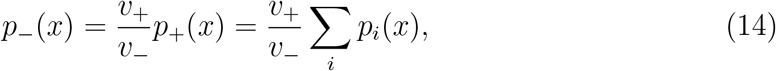

we can eliminate *p*_−_(*x*) in the stationary FPEs and achieve

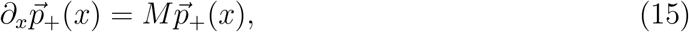

with 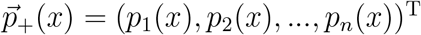, a matrix

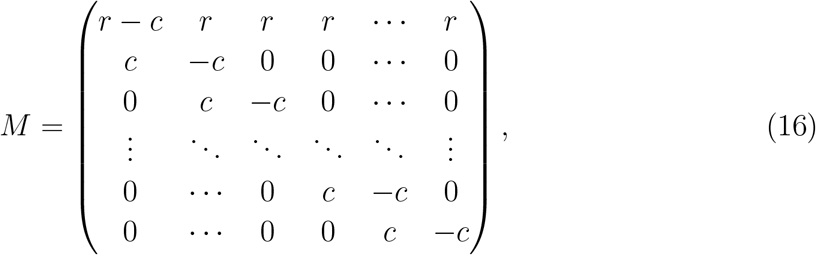

and the abbreviations *r* = *ω*_r_*/v*_−_ and *c* = *ω/v*_+_. Due to the reflecting boundary condition, a growing MT with length 0 must be in state 1, which provides the initial condition for the FPEs (15):

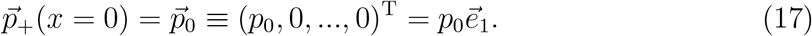

Therewith, the solution of the FPE (15) can formally be expressed as 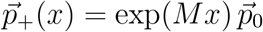. The parameter *p*_0_ is determined by the normalization condition 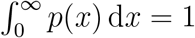, with

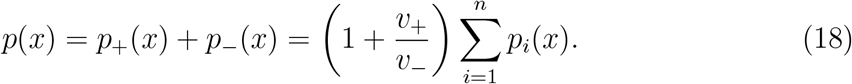

### 2.3. Force dependent growth velocities and catastrophe step rates

Here, we introduce force dependencies of the growth velocity and the catastrophe rate as similarly done in [17]. The effective velocity *v*_+_ of a growing MT results from the attachment of tubulin dimers to the MT tip with rate *ω*_on_ and their detachment with rate *ω*_off_ :

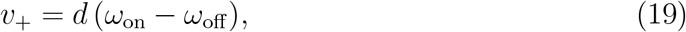

where *d* = 8 nm*/*13 ≈ 0.6 nm is the effective dimer size. An opposing force *F* modifies the attachment rate by a Boltzmann factor [19], yielding the force-dependent growth velocity

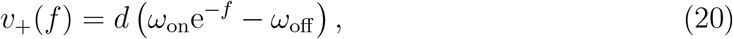

where we introduced the dimensionless force *f* = *F/F*_0_ with *F*_0_ = *k*_B_*T/d* ≈ 7 pN and the thermal energy *k*_B_*T* = 4.1 pN nm. As a consequence, MT growth is stalled (*v*_+_(*f*) = 0) at the dimensionless stall force

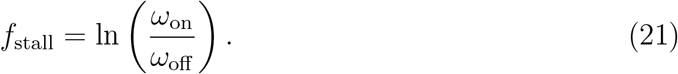

In the classical picture of dynamic instability, a growing MT is stabilized by a GTP-cap that is formed by GTP-tubulin dimers continuously added to the MT tip [1]. The GTP-tubulin dimers tend to be hydrolyzed after incorporation into the MT lattice so that the GTP cap may vanish and the MT undergoes a catastrophe. Since this is less likely if the addition of new GTP-tubulin dimers is fast, the catastrophe rate is expected to be negatively correlated with the growth velocity. Different microscopic models of dynamic instability were proposed that may be employed to deduce the velocity-dependence of the catastrophe rate [20, 21, 8, 9]. Here, we use the phenomenological approach from [17] that is based on the experimental observation that the mean growth duration ⟨*τ*_+_⟩ is a linear function of growth velocity,

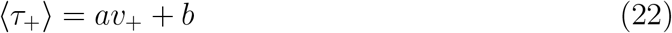

with *a* = 13.8 s^2^ nm^−1^ and *b* = 20 s [22].

For different multistep catastrophe mechanisms that yield the same experimentally observed average growth duration ⟨*τ*_+_⟩ = *nω*^−1^ but feature a different number of sub-states *n* (scenario (i)), this implies

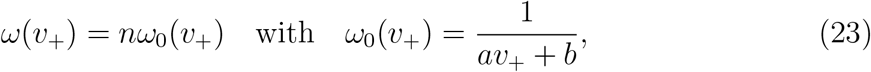

which conforms with (22) independently of the choice of *n*. Therefore, the step rate *ω*(*v*_+_) inherits a force dependence from the growth velocity via (20). Because the experimental observations suggest sub-steps that occur with the same rate *ω*(*v*_+_), also their force-dependence should be identical. Moreover, we assume that the number of catastrophe steps *n* is a structural feature of the MT that depends neither on the growth velocity nor on the applied force. Then, we find the force-dependent step rate

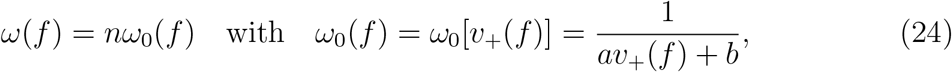

with *v*_+_(*f*) from (20). Analogously, we define *c*(*f*) = *ω*(*f*)*/v*_+_(*f*) and *c*_0_(*f*) = *ω*_0_(*f*)*/v*_+_(*f*).

### 2.4. Confinement between rigid walls

Above, we assumed unrestricted MT growth. In the following, we introduce a rigid wall that confines the MT length to a box between the reflecting boundary at *x* = 0 and the position of the wall *x* = *L*. Similar models for confined MTs with a single-step catastrophe were discussed in [16, 17]. If a MT tip reaches the wall, its growth is stalled (*v*_+_ = 0). A MT that reaches the wall in growing state *i* has to pass all *n* − *i* subsequent states to undergo a catastrophe and to leave the wall. Due to the velocity dependence of the catastrophe step rate (23), reaching the wall induces catastrophes with an increased step rate *ω*_L_ ≡ *ω*(*v*_+_ = 0).

We introduce probabilities *Q*_*i*_ to find the MT stalled at the wall while it is in growing state *i*. Since the MT leaves the wall instantaneously after a catastrophe, we can assume that *Q*_−_ = 0. The time evolution of *Q*_*i*_(*t*) is given by

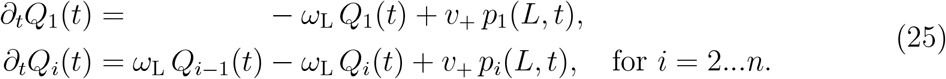

While the first terms describe the catastrophe steps of a MT that is already stalled, the expressions *v*_+_ *p*_*i*_(*L, t*) provide the probability currents onto the wall of a MT in state *i*. The total probability to find a MT tip at the wall is given by *Q*(*t*) ≡Σ_*i*_ *Q*_*i*_(*t*) and ∂_*t*_*Q*(*t*) = −*Q*_*n*_(*t*) + *v*_+_ *p*_+_(*L, t*) ≡ *J*_*L*_(*t*) yields the total probability current onto the wall. In the stationary state (∂_*t*_*Q*_*i*_(*t*) = 0), (25) is solved by

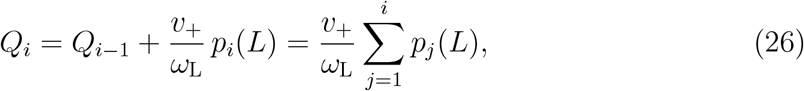

which adds up to the total probability

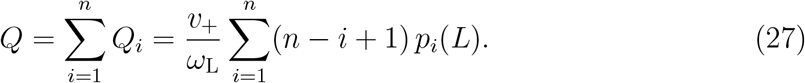

In the interior of the confining box (*x* < *L*), MT dynamics is still described by the FPEs (11). Therefore, the stationary probability density of the length of a confined MT is given by *p*_conf_ (*x*) = *p*_+_(*x*) + *p*_−_(*x*) + *Qδ*(*L* − *x*) and has to satisfy the normalization condition

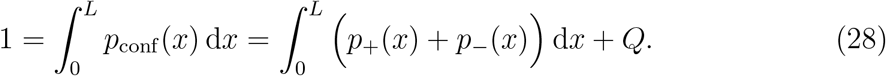

Since *Q* is a linear combination of *p*_*i*_(*L*), both *Q* and *p*_*±*_(*x*) are proportional to the constant of integration *p*_0_ introduced in the initial condition (17). Therefore, normalization can be achieved by adjusting *p*_0_, just like in the unconfined case.

## 3. Results

### 3.1. Bounded growth

We start our investigation of the multistep MT model with the force-free case. In general, the solution 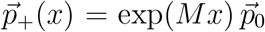 of the FPEs (15) can only be evaluated numerically, e.g., by numerical diagonalization of the coefficient matrix *M* as described in Appendix A. However, for the experimentally relevant case [6] that a MT can not be rescued (*ω*_r_ = *r* = 0) except for the forced rescue at the boundary *x* = 0, the solution can be expressed analytically:

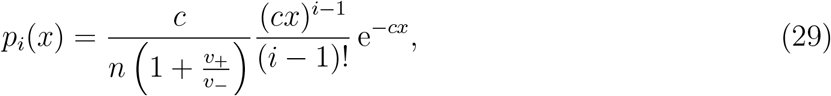

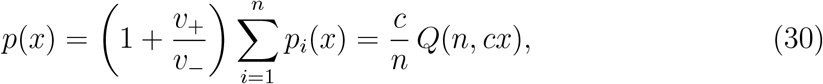

where *Q*(*n, x*) = Γ(*n, x*)*/*Γ(*n*) is the regularized form of the upper incomplete gamma function (see (8)).

In order to approach a solution of the general case (*ω*_r_ > 0), we make use of the results of Jemseena and Gopalakrishnan [15], who calculated the Laplace transform

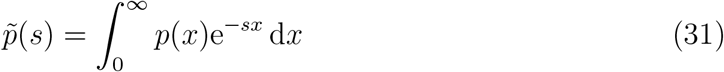

of the steady state length distribution for the case of an age-dependent catastrophe rate *ω*_c_(*τ*), where the age *τ* is the time that has passed since the last rescue event. Given an arbitrary probability density 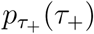 of growth durations *τ*_+_, the associated age-dependent catastrophe rate is given by (7). Combining that with the results of Jemseena and Gopalakrishnan [15], we find

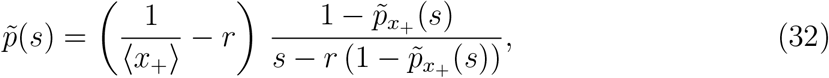

with the Laplace transform 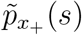 of the probability density of growth distances 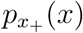. The derivation is presented in Appendix B. To achieve a result for the *n*-step catastrophe process, we substitute the Laplace transform of 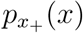 from (9) into (32):

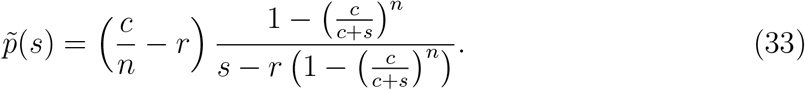

If *r* = 0, the inverse Laplace transform yields the probability density from (30). For the general case with rescues (*r* > 0), there is no analytical result for the inverse Laplace transform 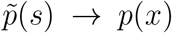. Nonetheless, we are able to compute exact results for the mean MT length ⟨*x*⟩ and the variance Var(*x*) = ⟨*x*^2^⟩ − ⟨*x*⟩^2^ by interpreting the Laplace transform as moment-generating function:

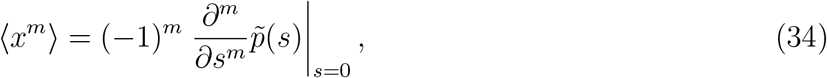

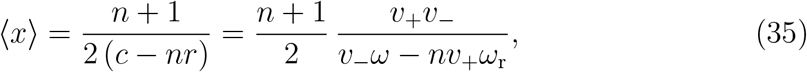

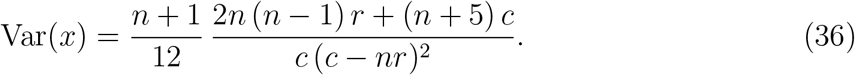

The method can be easily extended to higher moments and cumulants, which are listed in Appendix C up to the fourth degree.

Moments ⟨*x*^*m*^⟩ of the length distribution are dominated by large *x* and, thus, by the analytical properties of the Laplace transform 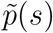 for small *s* around *s* = 0. On the other hand, we can approximate 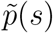 for large *s* as

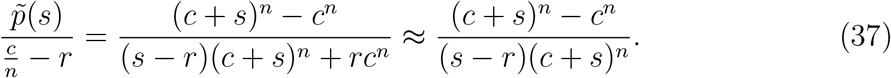

Then, the inverse Laplace transform is possible and provides an approximation of *p*(*x*) for short MT lengths:

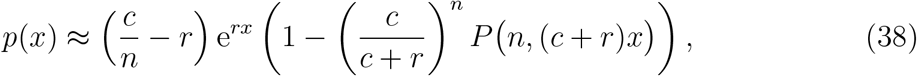

where *P* (*n, x*) = 1 − *Q*(*n, x*) is the regularized lower incomplete gamma function [18].

In the following, we compare our analytical results with stochastic simulations that solve the equation of motion of the MT for fixed time steps Δ*t* and include the random occurrence of a catastrophe step or a rescue after each time step. Here, we consider situation (i) and want to compare models with the same average growth duration ⟨*τ*_+_⟩ and, thus, the same parameter *V*_*n*_ in (10) but different number of sub-states *n*. Then, we have to use a catastrophe step rate proportional to *n*,

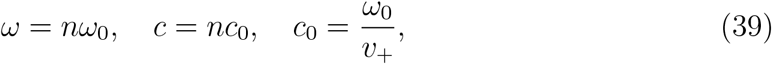

with a constant *ω*_0_, and the MT remains in the bounded regime (*V*_*n*_ < 0) independently of *n*. We use the following values:

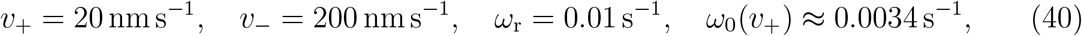

which are typical for bounded MT growth with negative *V*_*n*_ and correspond to tubulin concentrations around *c*_tub_ ∼ 10 μM ([17, 14] and references therein).

Figure 2 shows the results in absence of rescue events (*r* = 0) except for the forced rescues at *x* = 0. The analytical predictions from (29) and (30) perfectly match with the results from the simulations. The overall probability densities are monotonically decreasing functions and converge towards a step function 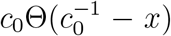 for large *n*. This uniform distribution for an infinite step catastrophe process can be made plausible by considering the growth distances *x*_+_: Since the standard deviation of the gamma distribution (9) is given by 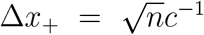, the relative error of growth distances 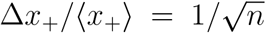 vanishes for large *n*. Moreover, as we assumed that *c* = *nc*_0_, also the absolute deviation decreases as 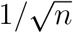 whereas the mean growth distance stays constant. Consequently, the more steps a MT has to pass to undergo a catastrophe, the more deterministic and predictable the length gain becomes. In the infinite step limit, the MT tip always covers the same distance during one growth interval. Then, in the absence of rescue events, a MT grows from *x* = 0 to exactly 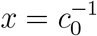 where it undergoes a catastrophe, and shrinks back to zero length where it is rescued again, finally resulting in a uniform distribution of MT lengths. Dynamically, repeated growth and shrinking by a sharp distance 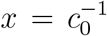 results in deterministic oscillations with a sharp period 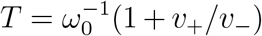. Similar effects have been discussed in [15]. Increasing the number of sub-states *n* at fixed mean growth duration thus sharpens the distribution of growth lengths and gives rise to quasi-deterministic MT length oscillations.

**Figure 2.**
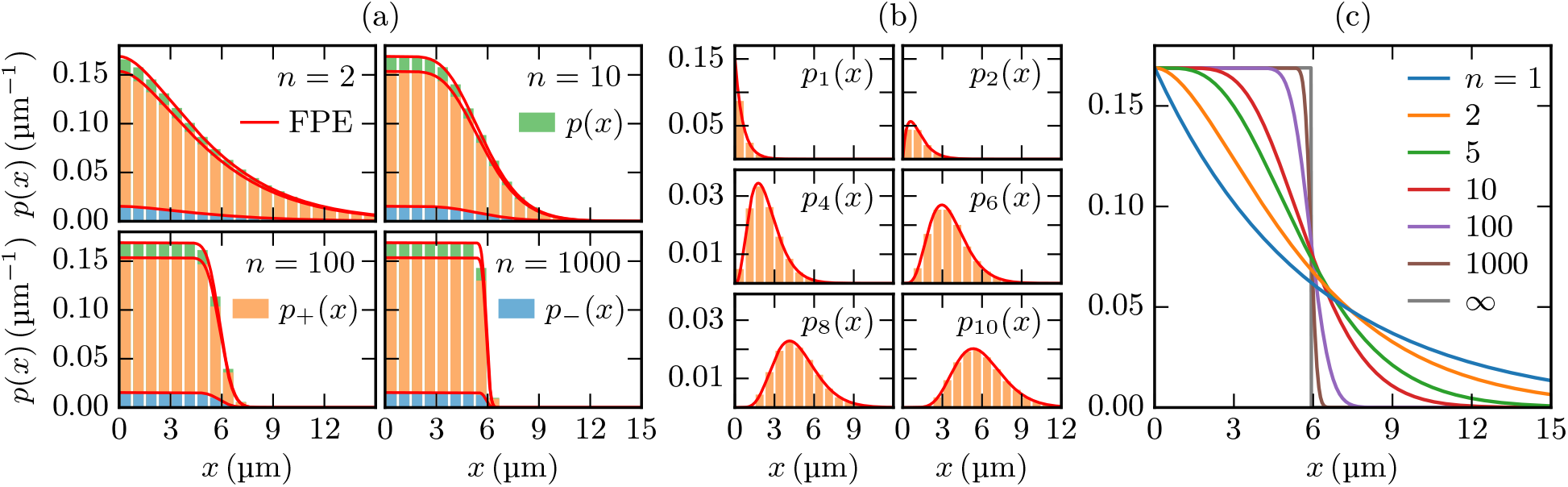
Comparison of analytical and simulation results in absence of rescue events. (a) Overall probability densities with various *n*. (b) Single state distributions in a 10-step process. For all cases in (a,b), the analytical FPE solutions (29) and (30) (red lines) match with the distributions measured in simulations (bars). (c) For an infinite-step process, the overall probability density converges to a step function.

If rescues are possible (*r* > 0), the probability density functions are not monotonic anymore but increase exponentially for short MT lengths up to a maximum, see figure 3(a). The exponential increase and the maximum are well described by the approximation (38). After the maximum, however, the approximation deviates from the real distribution. In that region, the probability densities measured in stochastic simulations are only fitted well by the numerical solution according to Appendix A, which is also the case for the single state densities *p*_*i*_(*x*) depicted in figure 3(b). If the number of steps *n* increases, the maximum becomes sharper and moves towards longer MT lengths up to 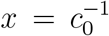. As we show in Appendix D, in the infinite step limit, the probability density approaches a piecewise defined function that initially grows exponentially as (*c* − *r*) exp(*rx*) until it has a step discontinuity at 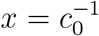. Moreover, there are non-analyticities of higher order at each multiple of 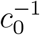. Like in the absence of rescues, this behavior can be explained with the determinism of MT growth distances in the infinite step limit: Since now the rescue rate is greater than zero, a MT can be rescued before shrinking to zero length and is able to grow beneath the single growth distance 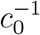. Nevertheless, a MT that grows from zero length after a forced rescue and undergoes a catastrophe at 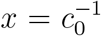 still shrinks back to zero with the probability exp(−*r/c*_0_) = 74 %. These 74 % alone would result in a step function again, and only in 74 % of the growth cycles that start from *x* = 0, the MT reaches lengths 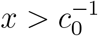, finally leading to the step discontinuity.

**Figure 3.**
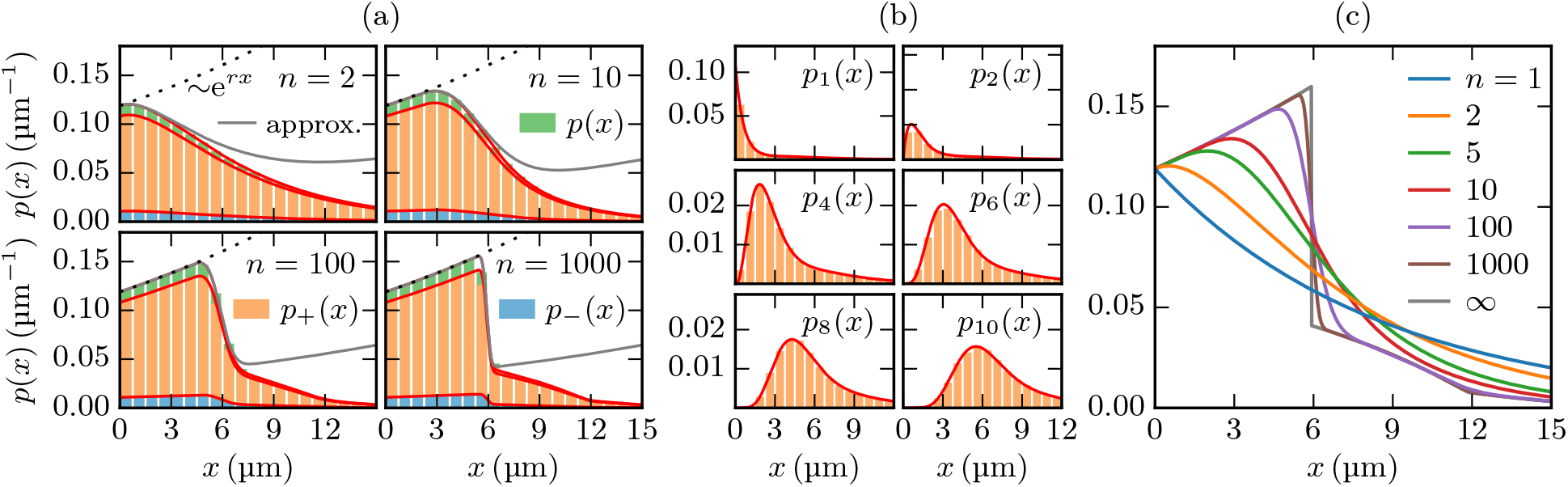
Comparison of deterministic and simulation results with *r* > 0. The deterministic results were calculated by numerically diagonalizing the matrix *M* from (16) as described in Appendix A. (a) Overall probability densities with various *n*. For short MT lengths, they grow exponentially (dashed lines) and can be approximated by (38) (gray lines). (b) Single state distributions in a 10-step process. For all cases in (a,b), the deterministic results (red lines) match with the distributions measured in simulations (bars). (c) For an infinite-step process, the overall probability density converges to a piecewise defined function that grows exponentially as (*c* − *r*) exp(*rx*) until it has a step discontinuity at 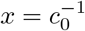, see Appendix D

As it can be seen in figure 4, (35) and (36) correctly describe the mean length and its variance as measured in stochastic simulations. If the number of catastrophe steps and the step rate are increased proportionally, the mean length decreases by up to one half of the single-step value.

**Figure 4.**
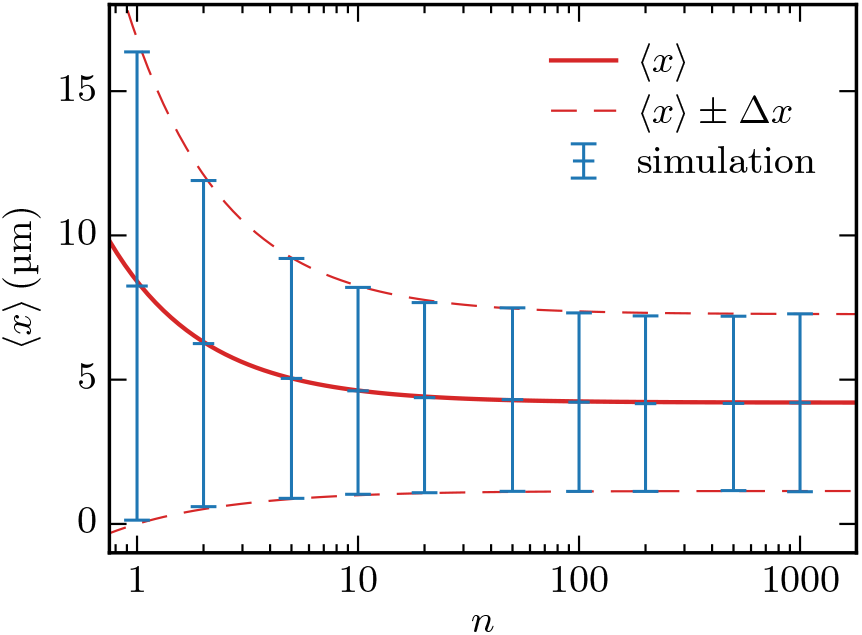
The mean MT length ⟨*x*⟩ and its standard deviation 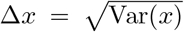 measured in simulations (blue) match with the analytical predictions (red) from (35) and (36).

The length distributions in Figs. 2(c) and 3(c) exhibit a characteristic change of shape as a function of the sub-state number *n*. Measurements of complete length distributions will, thus, allow to draw conclusions on this parameter related to the catastrophe mechanism. Also the reduction of the mean length ⟨*x*⟩ in Fig. 4 to one half of the single-step value for increasing *n* will allow to determine the sub-state number *n* if the average growth duration ⟨*τ*_+_⟩, rescue rate and growth and shrinkage velocities *v*_*±*_ are known.

### 3.2. Unbounded growth

In the regime of unbounded growth, there is no stationary solution which is why we have to analyze the time dependent FPEs (11). As described in detail in Appendix E, by use of the Fourier transform and the implicit function theorem, we are able to calculate a dispersion relation that is valid for long MTs. As *V*_*n*_ > 0 in the unbounded regime, the limit of long MTs is reached after long times. We conclude from the approximated dispersion relation that the probability density of MT lengths approaches a Gaussian distribution as in (4), but with the mean velocity *V*_*n*_ given by (10) and the diffusion constant

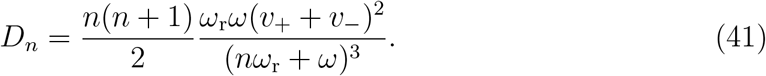

One can easily see that *D*_*n*_, and thus the standard deviation of MT lengths vanish for large *n* if the other parameters remain constant. Together with (10), this means that an increase of the number of catastrophe steps favors growth over shrinkage and thereby can make the MT leave the bounded regime, where a further increase of *n* favors growth even more until catastrophes are almost completely suppressed.

In figure 5, the results are compared to stochastic simulations. Here again, we consider situation (i) that the catastrophe step rate is proportional to *n* so that the mean growth duration and the mean velocity are constant. Then, the diffusion constant *D*_*n*_ does not vanish for large *n* but still decreases by up to one half of the single-step value. We use the same parameters as in (40) but with a ten times higher rescue rate in order to induce an unbounded state with *V*_*n*_ > 0. At the beginning of the simulations both the initial and, since the MTs are still short, the boundary condition cause the length distribution to significantly deviate from the Gaussian approximation. After long times, however, the MTs gained so much length that the probability of reaching the boundary is negligible. Then, the approximation provides an accurate description of the simulation results.

**Figure 5.**
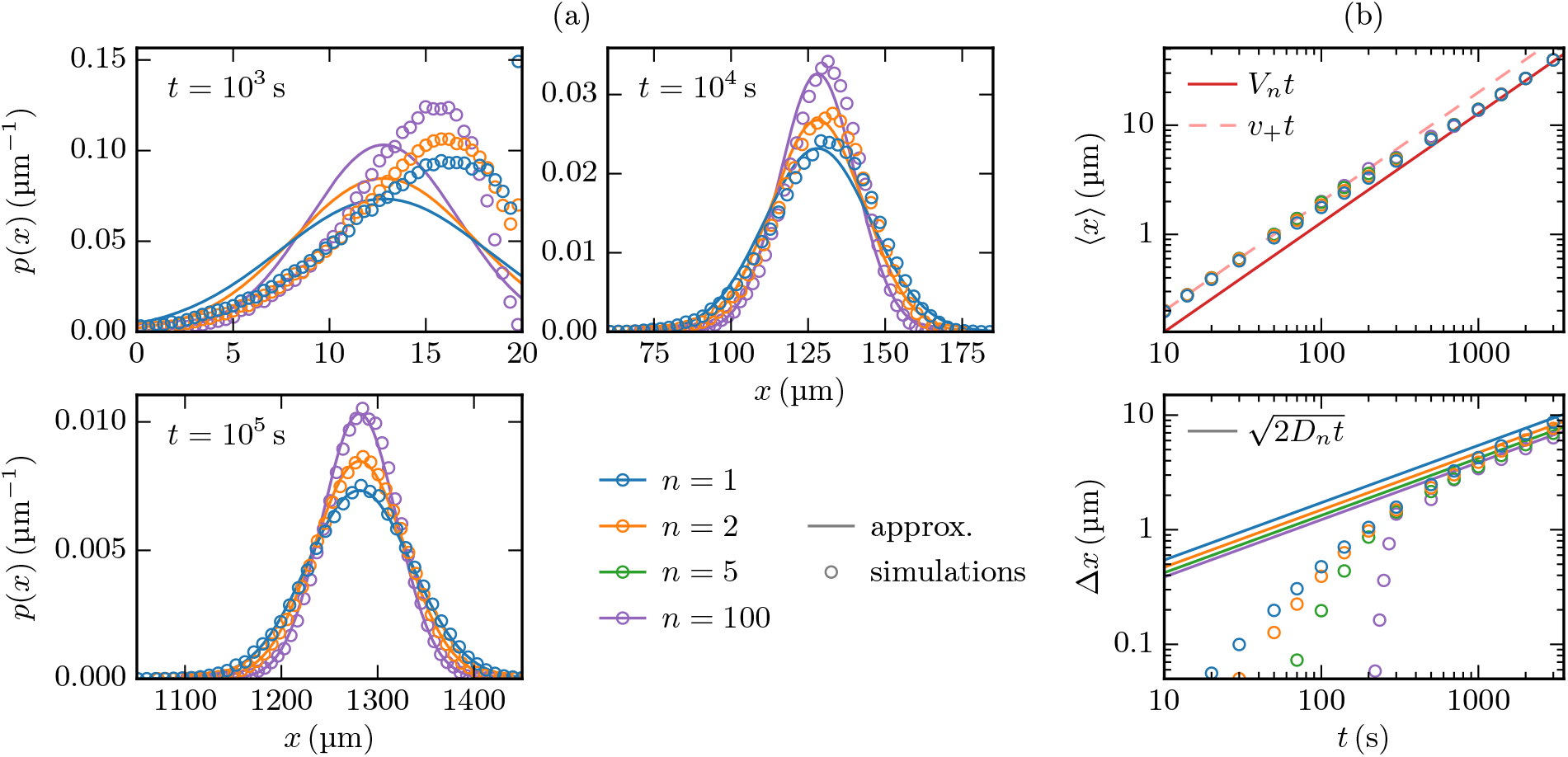
Comparison of analytical and simulation results for the unbounded case. The numerical results were each obtained from averaging over an ensemble of 10^5^ stochastic simulations of a MT that is initially in growing state 1 at *x* = 0. (a) Overall probability densities with various *n* at three different times, and the corresponding Gaussian approximations. The results only match for large times (*t* ≳ 10 000 s) whereas at the beginning (see *t* = 1000 s), the initial and the boundary condition, which are not incorporated in the approximation, still have a significant influence on the simulation results. For *n* = 1 and *n* = 2, there are even considerable amounts of MTs with length *v*_+_*t* = 20 μm that have not undergone a catastrophe yet (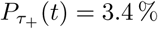 and 0.9 % respectively). (b) Time evolution of the mean MT length ⟨*x*⟩ and its standard deviation Δ*x* for various *n*. Until the first catastrophe, which occurs sharply at 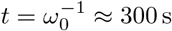 for *n* → ∞, the MT grows ballistically with ⟨*x*⟩ = *v*_+_*t* and Δ*x* = 0. Later, the simulations converge to the values predicted from the Gaussian approximation.

### 3.3. Microtubule growth against a force

An opposing force *f* affects the growth velocity and the step rate as described in section 2.3. We start our investigation with a MT that grows against a constant force and distinguish the two scenarios introduced above for the step rate: (i) We consider an experimentally given mean growth duration ⟨*τ*_+_⟩ as in (22), and consider models with different numbers *n* of sub-steps that conform with this experimental data. Then we use *ω*(*f*) = *nω*_0_(*f*) as in (24). (ii) Experimentally, also the number of sub-steps *n* might be changed *without* affecting the catastrophe step rate *ω*(*f*), e.g. by altering the MCAK concentration [6]. Then, we have a given *ω*(*f*), for which we use *ω*(*f*) = *ω*_0_(*f*) with *ω*_0_(*f*) from (24) to obtain agreement with the experimental data of [22] for *n* = 1.

For both scenarios, the opposing force influences the MT length distribution via a force dependent sub-step catastrophe rate *ω*(*f*) and via a force dependent growth velocity *v*_+_(*f*) as explained in section 2.3. Therefore, application of an opposing force does not give rise to novel shapes of MT length distributions but simply shifts parameters, in particular, the mean velocity parameter *V*_*n*_ = *V*_*n*_(*f*). This gives rise to other important effects, namely the existence of a critical force *f*_c_ for the boundary between bounded and unbounded regime [17].

While we vary *n, f* and *ω*_r_ in the following analysis, the other parameters are fixed throughout this section to

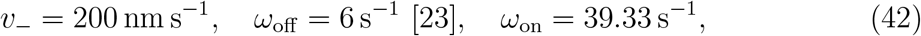

where *ω*_on_ was chosen such that the growth velocity and the step rate used in the previous sections (see (40)) are reproduced for *f* = 0.

Because of the force dependence of growth velocity and step rate, also the mean velocity *V*_*n*_ in (10) becomes force-dependent. In general *V*_*n*_ decreases with increasing force *f* such that a critical force *f*_c_ exists above which MT growth transitions from the unbounded to the bounded regime because it is suppressed by force. This critical force is determined from the condition *V*_*n*_ = 0 and is always smaller than the stall force *f*_stall_ = ln(*ω*_on_*/ω*_off_) = 1.88, which is set by the condition *v*_+_ = 0 so that the MT is not able to leave the boundary at *x* = 0 and, therefore, must be in the bounded regime.

For the first scenario (i) with a given ⟨*τ*_+_⟩ and *ω*(*f*) = *nω*_0_(*f*) as in (24), the velocity *V*_*n*_ is *n*-independent. Likewise, the critical force *f*_c_ is *n*-independent with

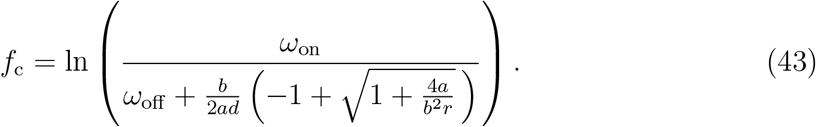

Therefore, measurements of the critical force will not allow to deduce information about the number of sub-steps *n*.

For the second scenario (ii), where the number of sub-steps *n* is changed without affecting the catastrophe step rate *ω*(*f*) = *ω*_0_(*f*) with *ω*_0_(*f*) from (24), on the other hand, the velocity *V*_*n*_ depends on *n* (see (10)). Therefore, we also find an *n*-dependent critical force

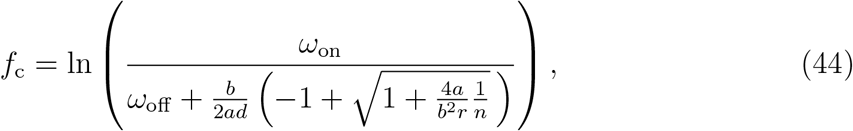

above which the MT is in the bounded regime. For both scenarios, the critical force is shown in figure 6(a) with different rescue rates *ω*_r_. In the second scenario, which applies to experiments employing MCAK [6], a value of *n* could be determined from experiments measuring the critical force *f*_c_. If the parameter values lie in the unbounded regime in the absence of force, the critical force *f*_c_ can be determined by increasing the force *f* such that MT growth becomes limited to a finite mean length ⟨*x*⟩ < ∞ for *f* = *f*_c_ at the transition to the bounded regime.

**Figure 6.**
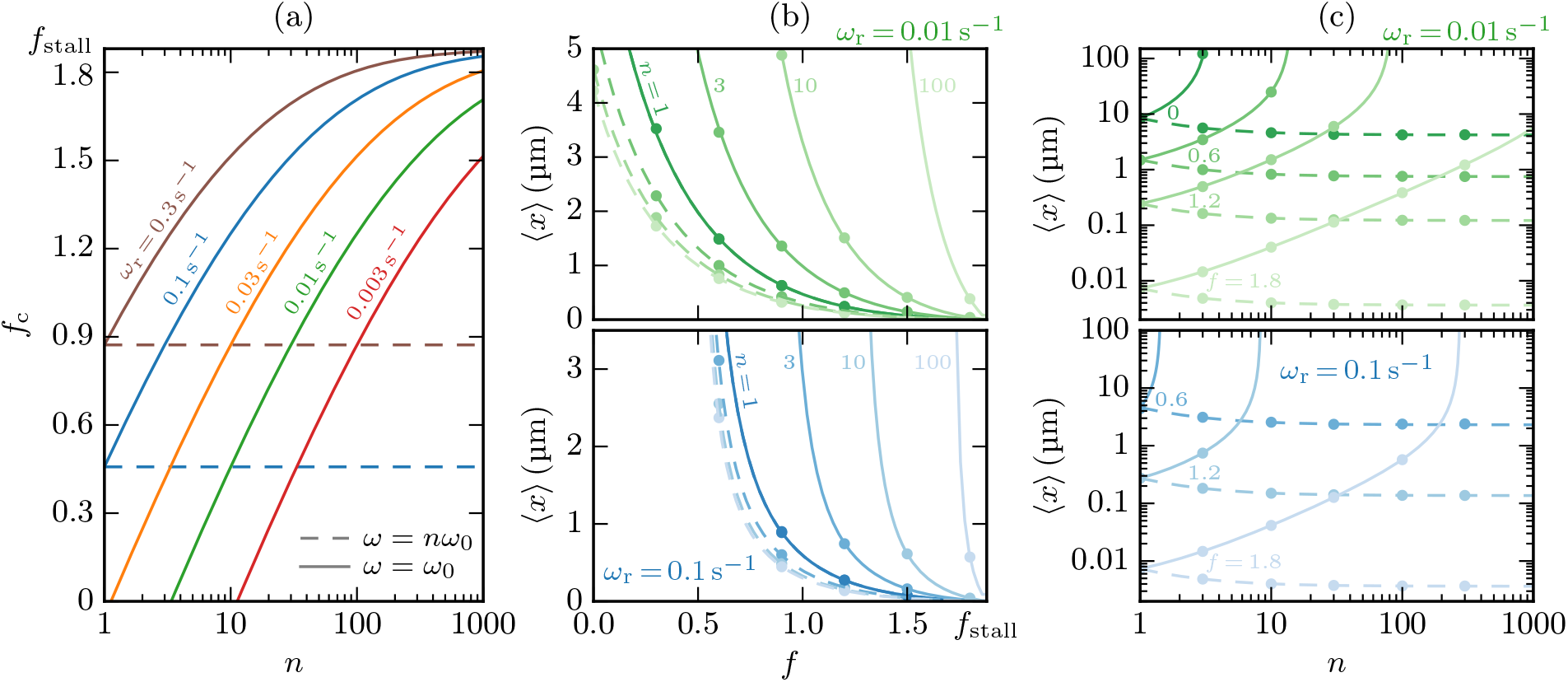
MT multistep dynamics under an opposing force *f*. The step rate is considered either as proportional to *n* (*ω*(*f*) = *nω*_0_(*f*), dashed lines) or as constant (*ω*(*f*) = *ω*_0_(*f*), solid lines). The dots represent results from stochastic simulations. The asymptotic behavior of the depicted plots is summarized in table 1. (a) Critical force *f*_c_ according to (43) and (44) as function of *n* for different rescue rates *ω*_r_. Above *f*_c_, MT growth is bounded, otherwise, it is unbounded. For *ω*(*f*) = *nω*_0_(*f*), *f*_c_ is constant and *f*_c_ < 0 if *ω*_r_ ≲ 0.03 s^−1^. For *ω*(*f*) = *nω*_0_(*f*), *f*_c_ converges to *f*_stall_ for large *n*. (b) Mean MT length ⟨*x*⟩ as function of the dimensionless force *f* for *n* = 1, 3, 10, 100 and *ω*_r_ = 0.01 s^−1^ (top) and *ω*_r_ = 0.1 s^−1^ (bottom). For both step rate scenarios, the opposing force reduces the mean MT length, which becomes 0 at *f* = *f*_stall_ and diverges at *f* = *f*_c_. (c) Mean MT length ⟨*x*⟩ as function of *n* for *f* = 0, 0.6, 1.2, 1.8 and *ω*_r_ = 0.01 s^−1^ (top) and *ω*_r_ = 0.1 s^−1^ (bottom). For *ω*(*f*) = *nω*_0_(*f*), ⟨*x*⟩ behaves as in figure 4 if the MT is in the bounded regime. For *ω*(*f*) = *ω*_0_(*f*), ⟨*x*⟩ increases with *n* and diverges at *n* = *n*_c_ = *ω*(*f*)*v*_−_*/ω*_r_*v*_+_(*f*).

Substituting *v*_+_ and *ω* in (35) with their force-dependent versions, we determine force dependent mean lengths ⟨*x*⟩, which are shown as function of *f* and *n* in figures 6(b) and (c), respectively. For the first scenario, we find a very weak *n*-dependence as in the absence of force (see (35) and figure 4). In the second scenario, where the number of sub-steps *n* is changed without affecting the catastrophe step rate, we find a much more pronounced *n*-dependence

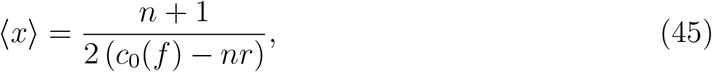

as also evidenced in figures 6(b) and (c). Again, we arrive at the conclusion that in the second scenario, a value for the sub-step number *n* could be determined from experiments measuring the mean length ⟨*x*⟩(*f*).

**Table 1.**
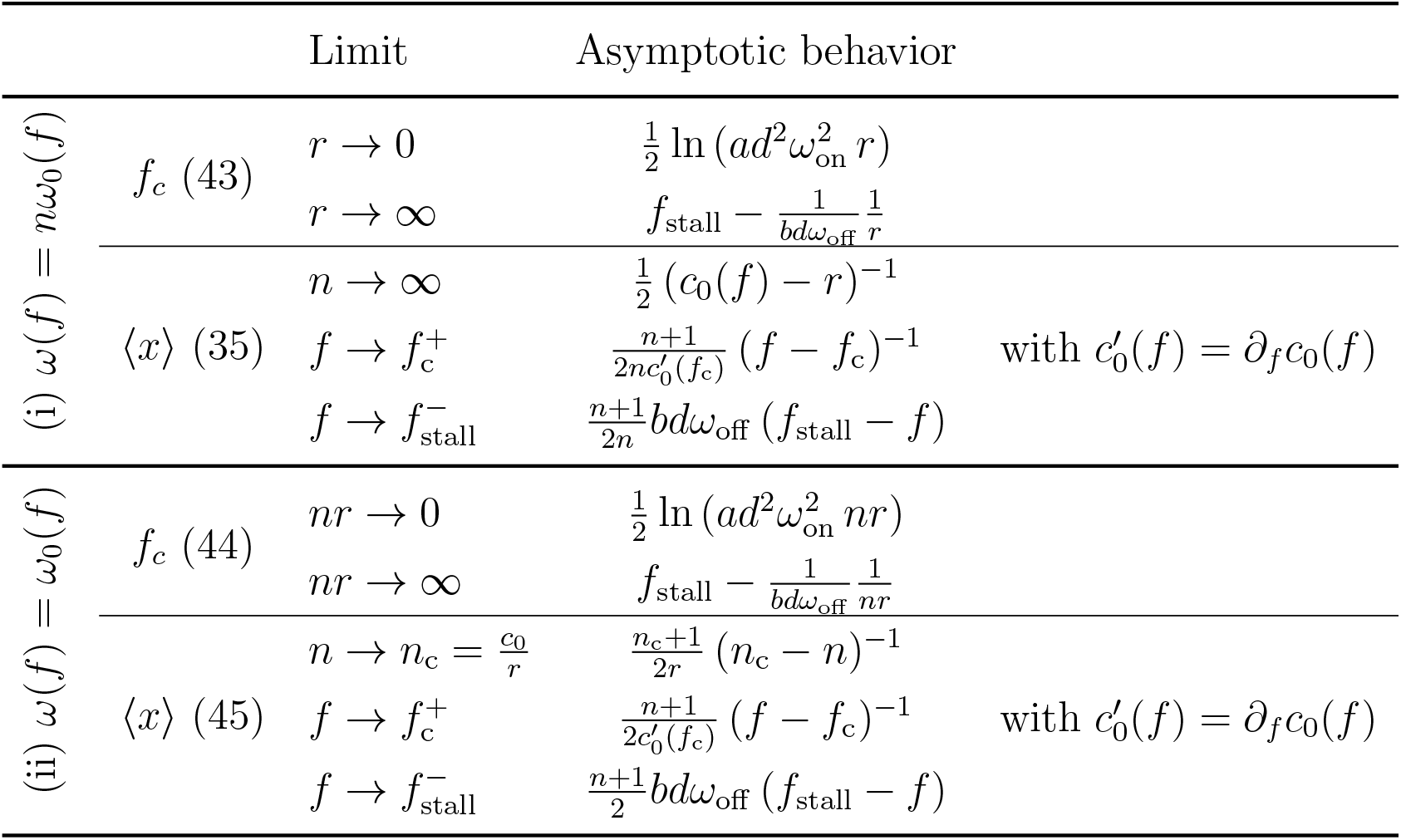
Asymptotic behavior of the critical force *f*_c_ and the mean MT length ⟨*x*⟩ in various limits, cf. figure 6.

The asymptotic behavior of *f*_c_ and ⟨*x*⟩ is summarized in table 1. In the second scenario, the critical force becomes a monotonic function of *n*, that grows logarithmically for small *n* and converges to *f*_stall_ for *n* → ∞, see figure 6(a) and table 1. Moreover, the mean length grows as a function of *n* and diverges at *n*_c_ = *ω*(*f*)*v*_−_*/ω*_r_*v*_+_(*f*) indicating a switch to the unbounded regime, see figure 6(c). Analogously, we observe diverging mean lengths in figure 6(b) for both scenarios when the applied force approaches the critical force *f*_c_.

### 3.4. Microtubule dynamics under a rigid confinement

So far, we assumed that the MT can grow freely to arbitrary lengths. However, the more realistic situation both in vitro and in vivo is that the MT is confined to a finite length. In the following, we investigate how multistep MT dynamics is affected by a rigid wall that restricts the MT length to *x* ≤ *L* as described in section 2.4. In contrast to the reflecting boundary at *x* = 0, the MT tip stays at the wall after reaching the length *L* until it has completed the remaining catastrophe steps with an increased step rate *ω*_L_ = *ω*(*v*_+_ = 0). In the stationary state, the probability *Q* to find the MT stalled at the wall (*x* = *L*) is given by (27) while the probability density in the interior (0 < *x* < *L*) is still defined by the FPEs (15). Therefore, the probability density *p*_conf_ (*x*) of a confined MT follows from truncating the unconfined probability density at *x* = *L* and weighting it with 1−*Q* in order to fulfill the normalization condition (28), see figure 7(a). As for an unconfined MT, the FPEs have to be solved numerically in the general case with rescues (*ω*_r_, *r* > 0), see Appendix A. However, in presence of a confinement to 0 < *x* < *L*, an analytical solution for moments is no longer possible via the Laplace transform because the Laplace transform is defined over the whole half-space 0 < *x*.

**Figure 7.**
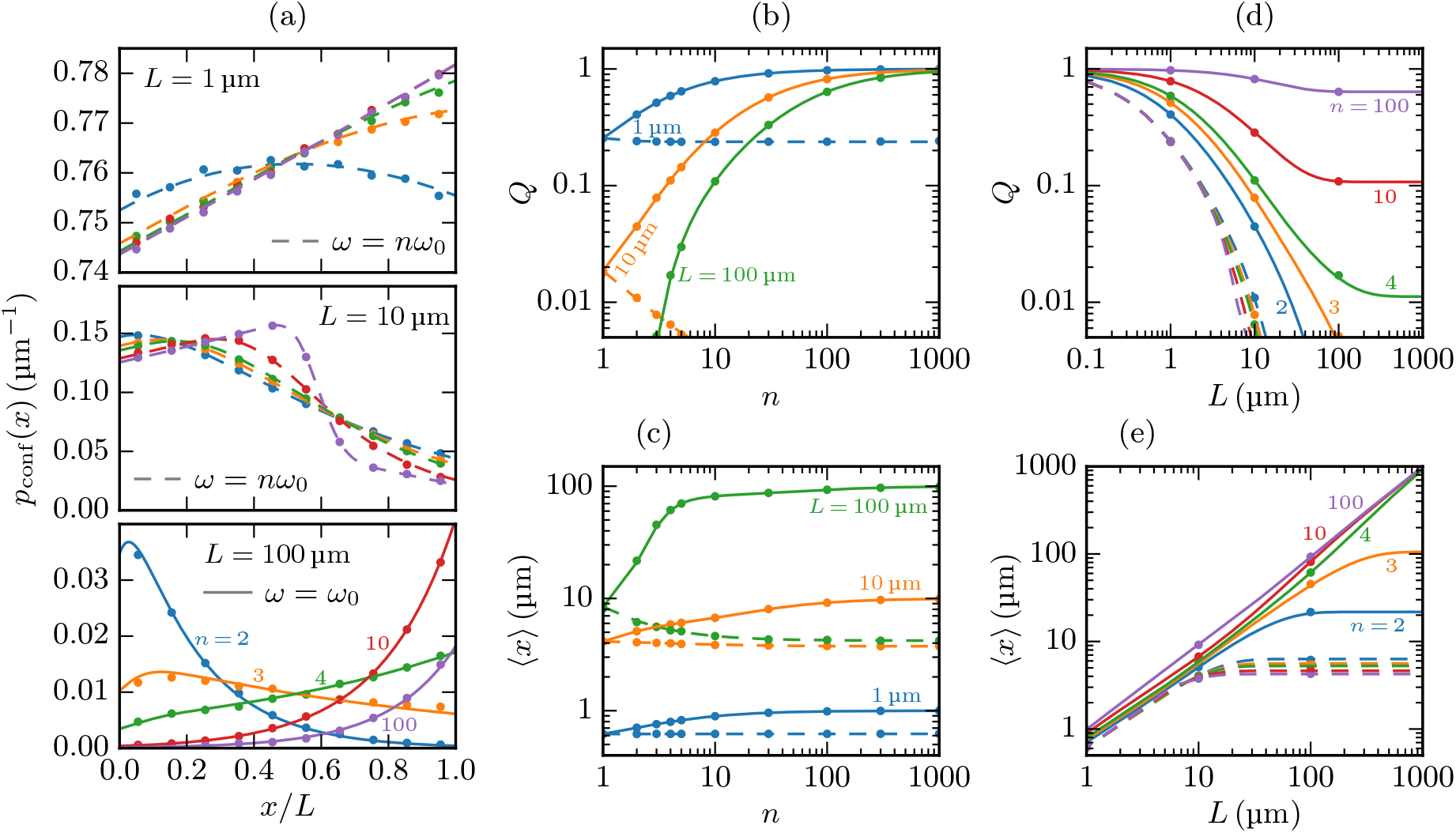
MT multistep dynamics confined by a rigid wall at *x* = *L*. Data is shown for *ω*_r_ = 0.01 s^−1^ and *f* = 0, i.e., for the parameters from (40). The step rate is considered either as proportional to *n* (*ω* = *nω*_0_, dashed lines) or as constant (*ω* = *ω*_0_, solid lines). The dots represent results from stochastic simulations. (a) Probability densities *p*_conf_ (*x*) in the interior of the box 0 < *x* < *L* for *n* = 2, 3, 4, 10, 100. The length distributions of a free MT as depicted in figure 3 are truncated at *x* = *L*. For *ω* = *nω*_0_ (top and center), the decisive factor for the shape of the distribution is whether the right boundary lies beyond the maximum, which coincides with the mean growth length 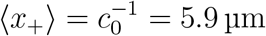 for *n* → ∞. For *ω* = *ω*_0_ (bottom), the MT switches from the bounded to the unbounded regime between *n* = 3 and *n* = 4 (*n*_c_ = 3.4). Accordingly, the MT lengths are shifted from being cumulated near the left boundary towards the right one. The total weight in the interior of the box decreases from *n* = 10 to *n* = 100 and further as the MT tends to be stalled at the wall for a longer time, see (b). (b,c) Probability *Q* to find the MT at the wall and mean MT length ⟨*x*⟩ as function of *n* for *L* = 1, 10, 100 μm. For *ω* = *nω*_0_, *Q* decreases with *n* due to the lighter tails of the length distributions. Since the MT is always in the bounded regime, the wall at *x* = *L* becomes irrelevant for *L* → ∞ and the mean length converges to (35), which is depicted in figure 4. For *ω* = *ω*_0_, a larger *n* shifts the MT length towards the right boundary and increases the residence time at the wall. Consequently, *Q* approaches 1 and ⟨*x*⟩ approaches *L* for *n* → ∞. (d,e) Probability *Q* to find the MT at the wall and mean MT length ⟨*x*⟩ as function of *L* for *n* = 2, 3, 4, 10, 100. If the MT is in the bounded regime (*ω* = *nω*_0_ or *ω* = *ω*_0_ and *n* = 2, 3), the wall can be neglected for large *L* so that *Q* vanishes and ⟨*x*⟩ approaches the mean length of a free MT from (35). In the unbounded regime (*ω* = *ω*_0_ and *n* = 4, 10, 100) the MT tends to the wall and is unlikely to shrink to zero length if *L* is large. Therefore, *Q* converges towards a finite value and ⟨*x*⟩ grows linearly with *L*.

Again, we distinguish the two scenarios (i) of a step rate that is proportional to *n* (*ω* = *nω*_0_) and (ii) of a constant step rate (*ω* = *ω*_0_), applying the force-free parameters from (40). Then, according to (23), the step rate at the wall is *ω*_L_ = *nω*_0_(*v*_+_ = 0) = *n/b* or *ω*_L_ = *ω*_0_(*v*_+_ = 0) = 1*/b*, respectively. In the proportional scenario (i), where the MT stays in the bounded regime for any *n*, the length distribution only retains the characteristics from figure 3 with a maximum and a sharp decrease thereafter if the position *L* of the wall lies beyond the mean growth length 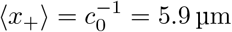, see figure 7(a). If *L* falls below the mean growth and the mean shrinkage length ⟨*x*_−_⟩ = *r*^−1^ = 20 μm, the MT length will approach a roughly uniform distribution as the MT is likely to both grow and shrink from boundary to boundary without catastrophes or rescues in the interior.

For a constant catastrophe step rate (ii), the MT switches from the bounded to the unbounded regime at *n* = *n*_c_ = 3.4. The growth of a MT in the unbounded regime is now confined by the rigid wall so that the MT lengths approach a stationary distribution as in the bounded regime. Since *V*_*n*_ > 0 in the formerly unbounded regime, the MT tends to the wall at *x* = *L* and greater MT lengths are more likely, see figure 7(a). It is important to note that the multistep characteristics of the length distribution, in particular the maximum, are lost if *V*_*n*_ > 0. Instead we find exponentially increasing probability densities because the distance from the right boundary that can be reached during a shrinkage-growth-cycle that starts from the wall is dominated by the single-step rescue. In order to restore a distribution with a maximum, rescue had to be a multistep process.

The behavior of the length distributions also becomes manifest in the stall probabilities *Q* and the mean lengths ⟨*x*⟩ depicted in figures 7(b–e). In the proportional step rate scenario (i), *Q* decreases with *n* (figure 7(b)) since the lighter tailed distributions make it less likely for the MT to reach the wall. With a constant step rate (ii), on the other hand, *Q* approaches 1 for two reasons: firstly, leaving the bounded regime by increasing *n* drives the MT towards the wall; secondly, the residence time at the wall is longer if more steps are required to leave it. As a consequence of *Q* → 1, the mean MT length approaches *L* if *n* is increased with a constant step rate, see figure 7(c).

Figures 7(d) and (e) show *Q* and the mean MT length, respectively, as a function of *L. Q*(*L*) decreases monotonically starting from *Q*(0) = 1, where the wall coincides with the reflecting boundary at *x* = 0. The behavior for larger *L* depends on whether an unconfined MT with the same parameters would be in the bounded (*V*_*n*_ < 0) or in the unbounded regime (*V*_*n*_ > 0): In the bounded regime, it is unlikely that the MT reaches the wall if *L* is large, and, hence, *Q*(*L*) vanishes while the mean length converges to the value of an unconfined MT as given by (35). For *V*_*n*_ > 0, on the other hand, the MT stays always in the vicinity of the wall so that its mean length becomes a linear function of *L*, and *Q* converges to a finite value because the probability density behaves as if the left boundary did not exist. We draw the general conclusion that the left (*x* = 0) or the right boundary (*x* = *L*) can be neglected for *V*_*n*_ > 0 or *V*_*n*_ < 0, respectively, if *L* is sufficiently large.

The results in figure 7 suggest that, in scenario (i), it might be possible to determine the number of catastrophe sub-steps *n* from the length distribution if the MT is confined to sufficiently large compartments *L* ≳ 10 μm so that the results for an unconfined MT can be applied. Then, also the probabilities *Q* to find the MT at the wall display a characteristic *n*-dependence, however, its absolute values are to small for an accurate measurement. In scenario (ii), which applies to experiments employing MCAK [6], a value of *n* could be determined more easily from length distributions and *Q*-measurement but also from experiments measuring the mean length ⟨*x*⟩.

## 4. Discussion

Based on experimental results that characterize MT growth periods as gamma distributed and conclude that catastrophe is a multistep process [4, 6], we extended the empirical Dogterom–Leibler model [2, 3] in order to analyze the consequences a multistep catastrophe mechanism has for the distribution of MT lengths. The multistep process has two main effects on the growth durations of a MT, which also underlie the consequential changes in the length distributions: Firstly, if the number of catastrophe steps is increased while keeping the rate of a single step constant, the growth durations become longer and the MT may leave the bounded regime. Secondly, the growth periods and hence the length gain during one growth interval are less stochastic if more steps are necessary to trigger a catastrophe.

In the case of bounded growth, the stationary length distribution has a steep descent in the vicinity of the mean growth distance *n/c*. In absence of rescues, the steep descent follows an area where the probability density is only slowly decreasing and becomes nearly constant for large *n*. If rescues are allowed, the distribution is exponentially increasing and has a maximum before it decreases sharply. In both cases, the length distributions are lighter tailed than the exponential distribution resulting from a single-step catastrophe, i.e., a multistep catastrophe reduces the number of MTs that are longer than the mean growth length. As a consequence, the mean MT length decreases by up to one half the single-step value if the number of catastrophe steps and the step rate are increased proportionally (see figure 4).

If the average growth duration ⟨*τ*_+_⟩ is known, we show that measurements of the shape of the bounded length distributions (Figs. 2(c) and 3(c)) or the mean MT length will give access to the number of sub-states *n* that best describe the experimental data (scenario (i)).

We also discussed the situation where MT dynamics is altered by MT regulators that do not only affect the velocities or the transition rates but the number of catastrophe sub-steps [6] (scenario (ii)). We conclude that by such a regulation, a MT acquires more potent ways to adapt to special situations inside the cell: while altering the classical four parameters only adjusts the range of MT lengths, which stay exponentially distributed in the single-step case (as long as they do not leave the bounded regime), variation of the additional parameter *n* changes the shape of the length distribution. As similarly discussed by Gardner *et al*. [6], this could be beneficial during mitosis. For instance, during prometaphase, the steep descent in the length distribution can appropriately limit the area that is explored by MTs in order to fasten search-and-capture of chromosomes [24]. In metaphase, accumulation of MT lengths around the maximum of the distribution may support the precise positioning of chromosomes in the metaphase plate and the maintenance of spindle length.

Stationary length distributions that have a maximum for short MT lengths before they apparently decrease exponentially and are similar to the ones in figure 3 were measured in several experimental studies [25, 26, 27, 28, 29, 30]. Though some of these studies have already been cited as evidence for a multistep catastrophe mechanism [6, 31], there are different reasons why this interpretation is dubious. In contrast to our model with a fixed minus and a dynamic plus end, the in vitro studies in [25, 29] examined free MTs that could polymerize simultaneously at both ends. Therefore, the shape of the length distribution can be rationalized by a convolution of the respective exponential distributions at the plus and the minus end, which are both obeying single-step dynamics [29]. In [26], the deviation from an exponential distribution for short MT lengths is attributed to the image resolution being to low to detect very short MTs. The results in [27, 28, 30] seem to be more in line with our model, yet these publications do not provide a quantitative evaluation of the measured length distributions, which makes a valid conclusion difficult. Besides, [27, 28] are in vivo experiments so that additional effects from microtubule associated proteins or spatial restrictions are likely.

In the regime of unbounded growth, the MT lengths approach a Gaussian distribution as in the single-step case but with a reduced variance. In vivo, the stabilization of MT growth due to an increase of the number of catastrophe steps might help interphase MTs, which have been shown to be in the unbounded regime [2, 32], to reach the cell boundary. On the other hand, at the transition from interphase to mitosis, MT lengths are significantly reduced in order to prepare the mitotic spindle assembly [33, 28]. The restructuring of the MT array may be supported by a reduction of the number of catastrophe steps, which destabilizes the MTs and shifts them to the bounded regime. This hypothesis is supported by the observation of Gardner *et al*. [6] that MCAK, which plays a key role for the control of MT dynamics during mitosis [34], promotes catastrophes by reducing the required steps from *n* = 3 to *n* = 1 and simultaneously keeping the step rate *ω* constant.

We also added a force-dependence of growth velocity and step rate to the model and analyzed how force affects multistep MT dynamics in the scenario of a step rate that is proportional to the number of required steps (*ω*(*f*) = *nω*_0_(*f*), scenario (i)) as well as for a step rate that does not depend on *n* (*ω*(*f*) = *ω*_0_(*f*), scenario (ii)). If the opposing force exceeds the critical force *f*_c_, the mean velocity *V*_*n*_ changes its sign and the MT switches from the unbounded to the bounded regime.

With a step rate that is proportional to *n*, the mean velocity and, thus, the critical force are constant. This scenario is useful for determining the number of sub-states *n* if the mean growth duration ⟨*τ*_+_⟩ = *nω*_0_ is known. Because the critical force is strictly *n*-independent, measurements of the critical force will not allow to deduce information about the number of sub-steps *n*.

When the step rate is independent of *n*, the force *f* and an increase of *n* counteract each other because the force suppresses growth and enhances catastrophe steps, thereby decreasing the mean growth duration and length, while a larger *n* results in an increased mean growth duration ⟨*τ*_+_⟩ = *nω*_0_. Thus, the critical force is a monotonic function of *n* and converges to the stall force, at which the growth velocity vanishes. Manipulating the step number without affecting the step rate is possible by a variation of the MCAK concentration [6]. In this situation measurements of the critical force *f*_c_ give information on the value of *n*.

If the MT is confined to finite lengths *L*, for instance by the cell cortex, the MT will either tend to shrink back to zero length if it is in the bounded regime, or it will grow repeatedly against the confining boundary if MT growth would be unbounded without the confinement. In the latter case, the stationary length distribution has no maximum but increases exponentially up to the boundary. Since this is due to the single-step kinetics of the rescue, a maximum could be restored by a multistep rescue, which, however, has not been observed so far to our knowledge. As aforementioned, MTs are in the unbounded regime during interphase [2, 32], where the microtubule organizing center is positioned by direct pushing and/or by dynein-mediated pulling interactions between the plus ends and the actin cortex [35, 36, 37, 38]. Since it would be disadvantageous for these interactions if the MT distribution had a maximum and the most probable MT length lay in front of the cell cortex, a single-step rescue and an exponential length distribution are indeed favorable in this situation. Moreover, we argued above that interphase MTs might be in the unbounded regime as a result of an increased number of required catastrophe steps *n*. This mechanism would further support the interaction with the cell cortex since the probability *Q* to find the MT plus end at the confining boundary correlates positively with *n* if the step rate is constant.

## 5. Conclusion

In conclusion, a catastrophe mechanism that requires multiple steps has significant effects on the length distribution of MTs. Modifying the number of required catastrophe steps, e.g., by regulation via MCAK [6], allows to adapt not only the scale but also the shape of the length distribution to be beneficial for the present physiological situation. The multistep characteristics of MT length are retained under an opposing force; the critical force to suppress unbounded growth becomes sensitive to modification of the number of catastrophe steps. Confined MTs can continue to show characteristic maxima in the MT length distribution only in the bounded regime and if the compartment is sufficiently long; otherwise MT, length distributions tend to simple exponentially decreasing or increasing distributions in the bounded and unbounded case, respectively.

## Appendix A. Numerical solution of the stationary Fokker–Planck equations

We solve the stationary FPE (15) by numerical determination of the eigenvalues *λ*_*j*_ and -vectors 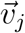 of the matrix *M*. The solution can then be written as

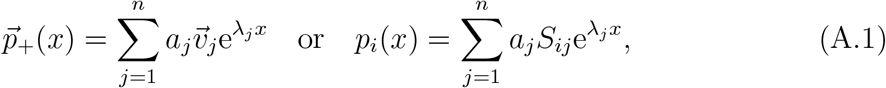

with 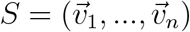. The coefficients *a*_*j*_ follow from the initial condition (17) at *x* = 0:

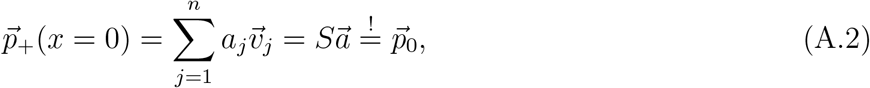

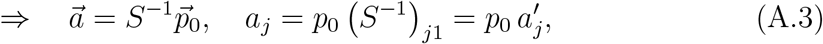

with 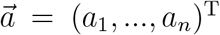. Finally, the constant *p*_0_ is determined by the normalization condition (note that Re *λ*_*j*_ < 0 for any eigenvalue):

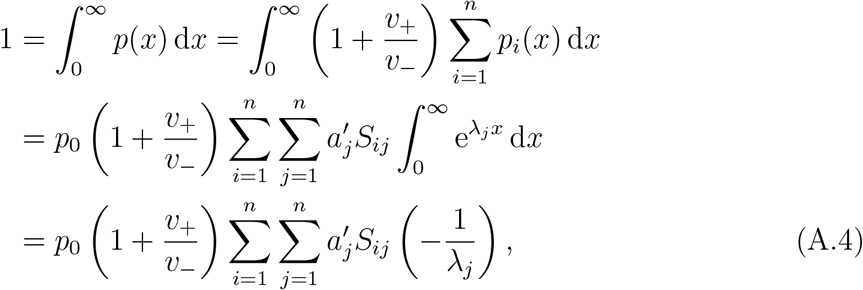

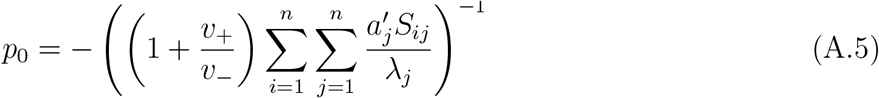

If a MT is confined to *x* ≤ *L* by a rigid wall as described in section 2.4, the probability density in the interior of the box is still given by (A.1) and (A.3), but is accompanied by the probability *Q* to find the MT tip stalled at the wall, which is a linear combination of *p*_*i*_(*L*) in the stationary state, see (27). Therefore, for a confined MT, the constant *p*_0_ follows from the normalization condition (28), which has to be fulfilled by the total probability density *p*_conf_ (*x*) = *p*(*x*) + *Qδ*(*L* − *x*):

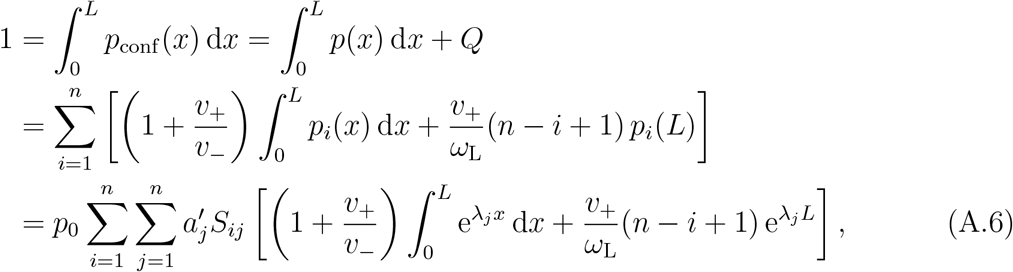

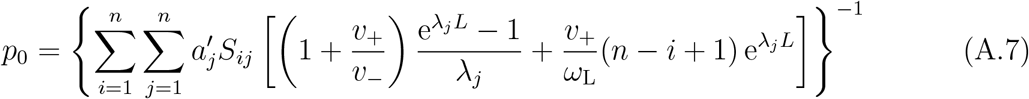

## Appendix B. Solution for an age-dependent catastrophe rate

Jemseena and Gopalakrishnan [15] found that in case of an age-dependent catastrophe rate *ω*_c_(*τ*) and a reflecting boundary at *x* = 0, the Laplace transformed overall probability density 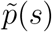 of MT lengths is given by

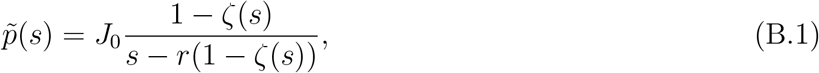

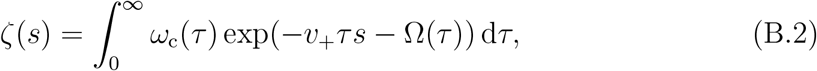

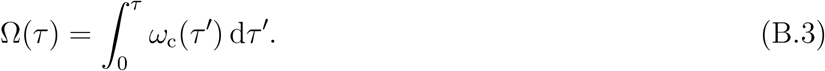

Using (34), the normalization condition, which defines the constant *J*_0_, can be expressed as

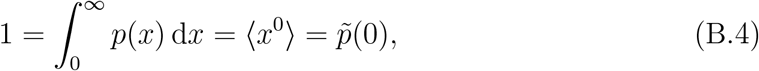

Substituting the catastrophe rate 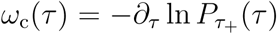 that follows from an arbitrary distribution of growth durations (see (7)), we find

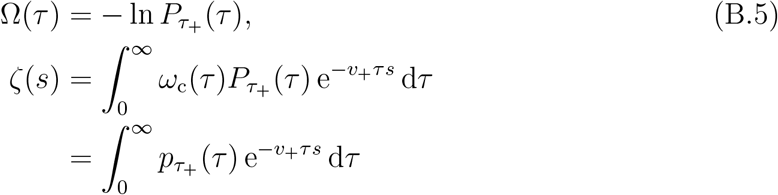

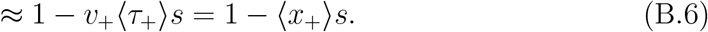

The last line is valid for small *s*. Next, we substitute *x* = *v*_+_*τ* to show that *ζ*(*s*) is the Laplace transform of the probability density of growth distances:

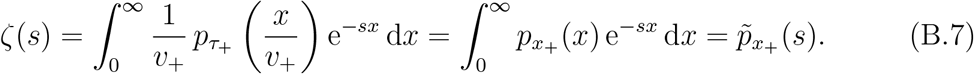

For the purpose of normalization, we use the approximated form of *ζ*(*s*) from (B.6):

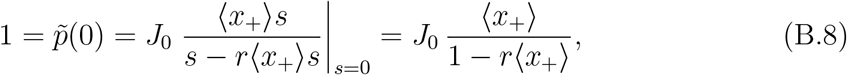

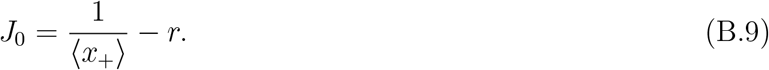

Substituting (B.7) and (B.9) into (B.1) finally results in 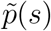 as given in (32).

## Appendix C. Moments and cumulants of MT length

The moments ⟨*x*^*m*^⟩ and the cumulants *κ*_*m*_ of MT length *x* can be derived from the Laplace transformed probability density in (33) as

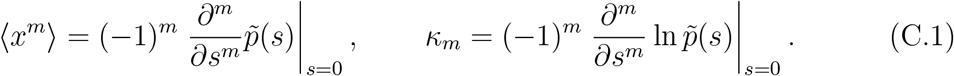

The first four moments and cumulants are:

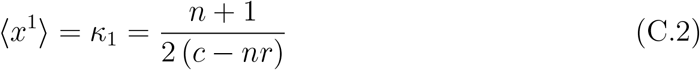

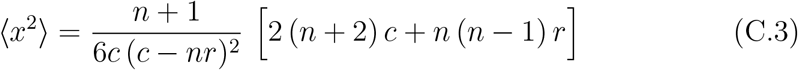

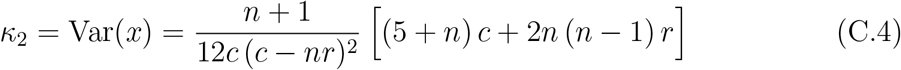

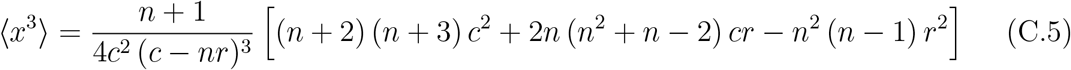

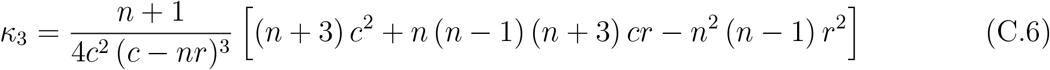

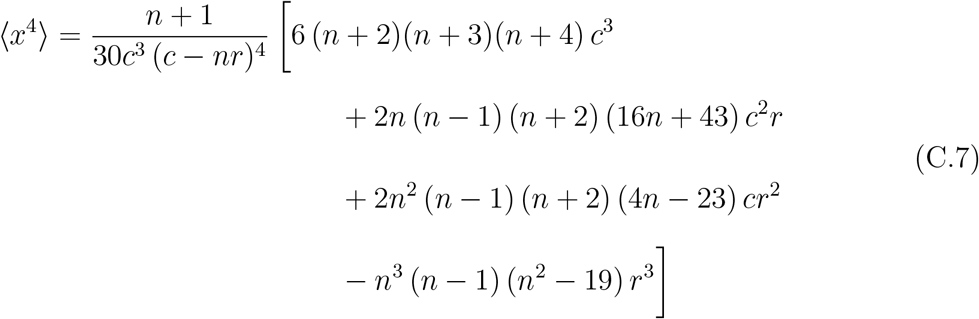

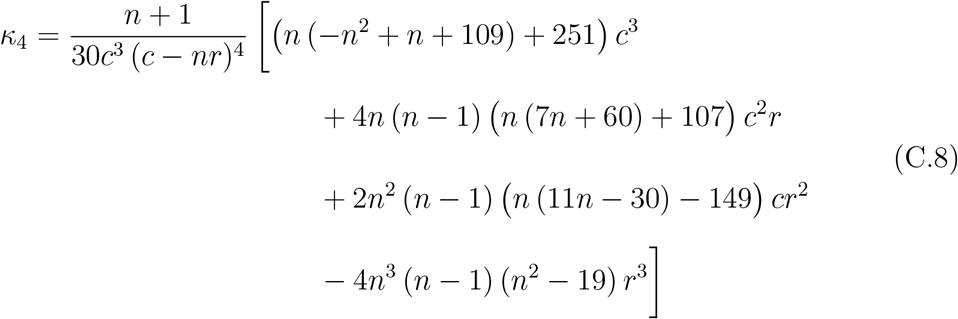

## Appendix D. Stationary solution for an infinite step catastrophe process

In the limit *n* → ∞, the Laplace transformed probability density of growth distances is 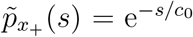. For the Laplace transform of the overall probability density of MT lengths in presence of rescue events follows:

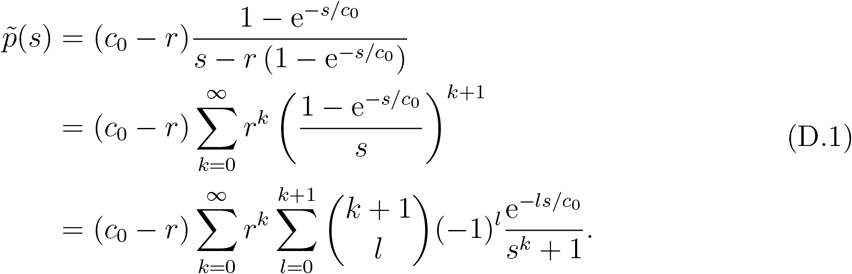

Inverse Laplace transformation yields:

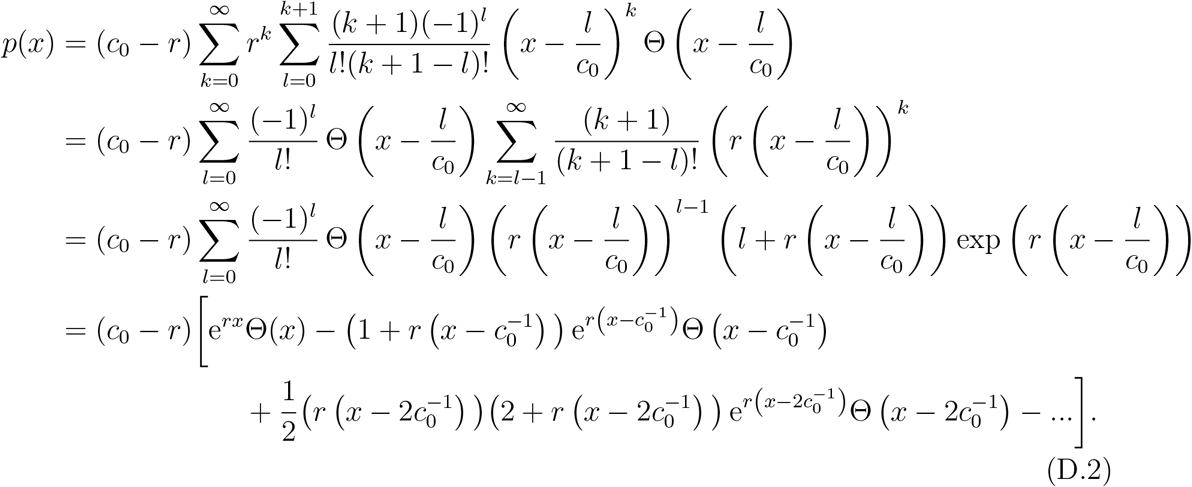

We see that the probability density increases exponentially until it has a step discontinuity at the (now deterministic) growth length 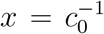. At each multiple of the growth length, the function is non-analytic since an additional term contributes thereafter. The first two non-analyticities can be seen in figure 3(c). In absence of rescues (*r* = 0) the probability density correctly turns into the step function 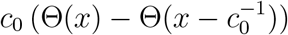.

## Appendix E. Approximation for unbounded growth

We define the Fourier transform

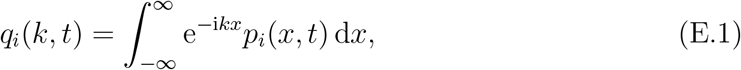

and apply it to the FPE (11):

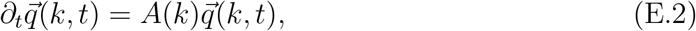

with 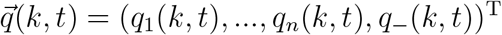 and

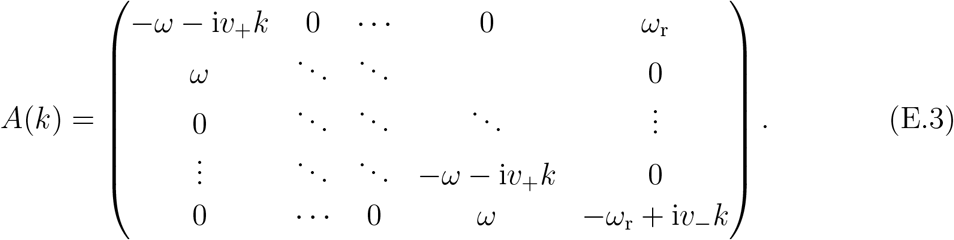

The fundamental solutions of (E.2) are given by

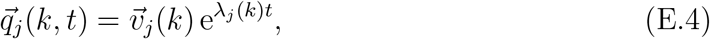

where *λ*_*j*_(*k*) and 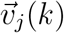 are the eigenvalues and eigenvectors of *A*(*k*), respectively. The eigenvalues can be determined from the characteristic polynomial of *A*(*k*):

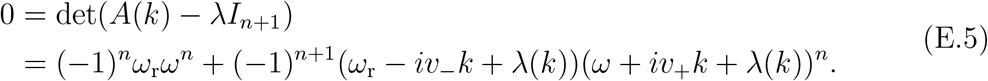

An approximation for long MTs in real space corresponds to the limit of small *k* in Fourier space. For *k* = 0, *λ*_0_(*k* = 0) = 0 is the only non-negative eigenvalue and, therefore, provides the only solution 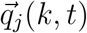 that does not vanish for *t* → ∞. To approximate the dispersion relation for long MTs after long times, we expand *λ*_0_(*k*) around *k* = 0 using the implicit function theorem. According to (E.5), *λ*_0_(*k*) can be implicitly defined by *f* (*λ*_0_(*k*), *k*) = 0 with

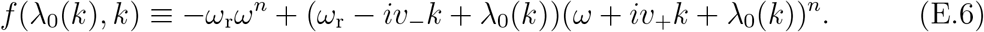

From the implicit function theorem and with *λ*_0_(*k* = 0) = 0, we find

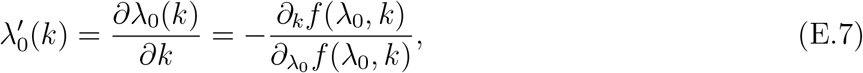

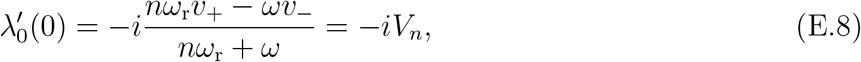

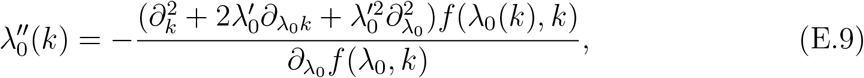

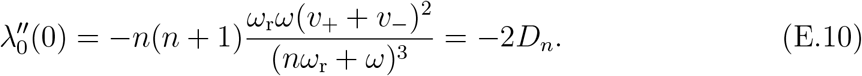

This results into the dispersion relation

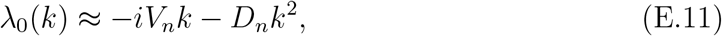

which corresponds to a diffusion process with diffusion constant *D*_*n*_ and drift *V*_*n*_, which is the mean velocity as deduced in (10).

Normalization of the overall probability density is translated into Fourier space by the condition *q*(0, *t*) = 1, where *q*(*k, t*) is the Fourier transform of the overall probability density *p*(*x, t*). Finally, an inverse Fourier transform of *q*(*k, t*) = exp(*λ*_0_(*k*)*t*) yields the approximation of *p*(*x, t*) for long times:

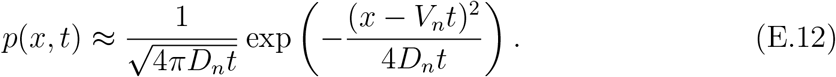

